# A scaffold attachment factor PHM-2 regulates synaptic transmission through SLO-2 potassium channel in *C. elegans*

**DOI:** 10.64898/2025.12.24.696350

**Authors:** Longgang Niu, Karthika Murugasen, Shannon Honggodo, Sakia Ferdousy, Lishuang Zhu, Bojun Chen

## Abstract

Scaffold attachment factor B (SAFB) proteins are evolutionarily conserved DNA/RNA binding proteins that are involved in multiple processes of gene expression. These proteins are broadly expressed with particular high expression observed in the nervous system. However, their physiological roles in neurons are largely unclear. Here we show that PHM-2, the sole SAFB ortholog in *C. elegans*, regulates synaptic transmission at the neuromuscular junctions through an effect on SLO-2 potassium channel. We found that *phm-2* knockout suppresses a sluggish phenotype of worms expressing a hyperactive SLO-2 channel, greatly reduces SLO-2-mediated neuronal whole-cell currents, and enhances neuromuscular synaptic transmission. In addition, we found that PHM-2 genetically interacts with another DNA/RNA binding protein, HRPU-2/hnRNP U, to control SLO-2 expression through a posttranscriptional mechanism. These results reveal a novel function of a SAFB protein in regulating neuronal activity, and may help understand the physiological roles of SAFB proteins in the nervous system of other species.

**Author Summary:** Proteins in the SAFB family are found in many species, and they help control how genes are expressed in cells. These proteins are commonly present in the nervous system, but their exact roles in nerve cells are not well understood. In this study, we examined the single SAFB-like protein called PHM-2 in the nematode *C. elegans* to learn how it affects the nervous system. We discovered that PHM-2 plays an important role in communication between nerve cells and muscles. Worms lacking PHM-2 were able to counteract the sluggish movement caused by a hyperactive potassium channel called SLO-2. Without PHM-2, nerve cells had much smaller electrical currents mediated by SLO-2 and a stronger signaling from nerves to muscles. We also found that PHM-2 works together with another genetic regulator, HRPU-2, to control the amount of SLO-2 protein made in nerve cells. These findings reveal a new role for SAFB proteins in shaping neuronal activity by regulating potassium channels. Understanding this process in worms may provide clues about how these proteins contribute to brain function in humans.

## Introduction

RNA-binding proteins (RBPs) constitute one of the largest protein families in eukaryotic cells. These proteins assemble with RNAs to form ribonucleoprotein complexes and play critical roles in RNA biogenesis, stability, function, transport, and cellular localization (1). In human cells, thousands of RBPs associate with their target RNAs and other proteins to form extensive regulatory networks that regulate cell homeostasis. Analysis of disease association data has identified over a thousand RBPs that are mutated in various human genetic diseases, and Gene Ontology analysis shows that mutations of RBPs are predominantly associated with metabolism and nervous system development (2). In the nervous system, where gene expression is highly dynamic during development and across different regions (3, 4), RBPs are essential to neurogenesis, differentiation, and synaptic plasticity, and deficiencies in their expression and/or distribution may cause neurologic disorders such as intellectual disabilities, motor impairments, and neurodegeneration (5, 6). Undoubtedly, mechanistic dissection of the specific roles of RBPs in the brain is crucial for understanding the molecular basis of neurological diseases.

The scaffold attachment factor B (SAFB) proteins are a group of evolutionarily conserved RBPs comprising three members including SAFB1, SAFB2, and SLTM (SAFB-like transcriptional modulator) (7). Studies have shown that SAFB proteins play important roles in many aspects of cellular processes, such as DNA repair (8, 9), cellular stress response (10-12), transcription (13-16), and processing of mRNA and miRNA (17-20). Evidence also suggests that SAFB proteins may be involved in the progression of various cancers, including prostate cancer (21), pancreatic adenocarcinoma (22), breast cancer (23), and bladder cancer (24). Like many other RBPs, SAFB proteins are abundantly expressed in the brain (7, 25). In cortical and hippocampal primary neuronal cultures, expression of the SAFB proteins is primarily found in the nuclei, with SAFB2 and SLTM also co-localized in the same dendritic puncta (7). Adenovirus-mediated expression of SAFB1 in primary hippocampal neurons results in increased dendritic spine size (20). SAFB1 expression and localization are abnormal in the post-mortem brain tissue of spinocerebellar ataxias and Huntington’s chorea patients (26). A recent study shows that SAFB interacts with the ribonuclease Drosha to regulate hippocampal stem cell fate (27). These observations suggest that SAFB proteins are important for neuronal function, but their physiological roles in the nervous system remain largely undefined.

Slo2 channels are large-conductance potassium channels present in mammals and invertebrates (28, 29) and are major contributors to delayed outward currents in neurons (30, 31). Humans and mice each have two Slo2 channel, Slick/Slo2.1 and Slack/Slo2.2, which are broadly expressed in the nervous system (32-34). These channels shape neuronal excitability and act as key suppressors of sensory hypersensitivity, seizures, and excitotoxicity (28) (35-37). Studies have shown that Slack regulates working memory through HCN coupling (38), supports hippocampal plasticity (39), and decreases amygdala excitability to limit anxiety (40). Slack also links neuronal activity to local translation via two key mRNA translation regulators, FMRP and CYFIP1 (41), and interacts with Na(V)1.6 to influence excitability of excitatory and inhibitory neurons (42). In this study, we found that neuronal expression of SLO-2, an ortholog of human Slo2.2/Slack in *C. elegans*, is regulated by the SAFB ortholog PHM-2. Loss of *phm-2* reduces SLO-2-dependent whole-cell currents in motor neurons and enhances synaptic transmission at the neuromuscular junction. We further show that PHM-2 cooperates with another RBP, HRPU-2/hnRNP U, to control SLO-2 expression in neurons. Our findings reveal a molecular mechanism by which a SAFB protein regulates neuronal function and may shed light on the physiological roles of SAFB proteins in the nervous system of other species.

## Results

### phm-2 knockout suppresses sluggish phenotype of slo-2(gf) worms

In an effort to identify novel regulators of the SLO-2 potassium channel in *C. elegans*, we performed a forward genetic screen for suppressors of a sluggish phenotype of worms expressing a hyperactive SLO-2 channel. The hyperactive or *gain-of-function* (*gf*) SLO-2 channel was engineered by mutating three consecutive amino acid residues, GQT, to AEL in the 6^th^ transmembrane domain of SLO-2 subunit (43).

Worms expressing the SLO-2(*gf*) channel showed greatly reduced overall locomotion speed and a reduced forward/backward ratio (**Figure 1A, B**). This phenotype was almost completely suppressed in one of the isolated mutants, *zw67*. Through an SNP-based mapping approach (44), we mapped the mutation to a small region on the right arm of chromosome I. Analysis of the whole-genome sequencing data identified a nonsense mutation in the gene *phm-2* (*F32B4.4*, www.wormbase.org) within that region.

**Figure 1.**
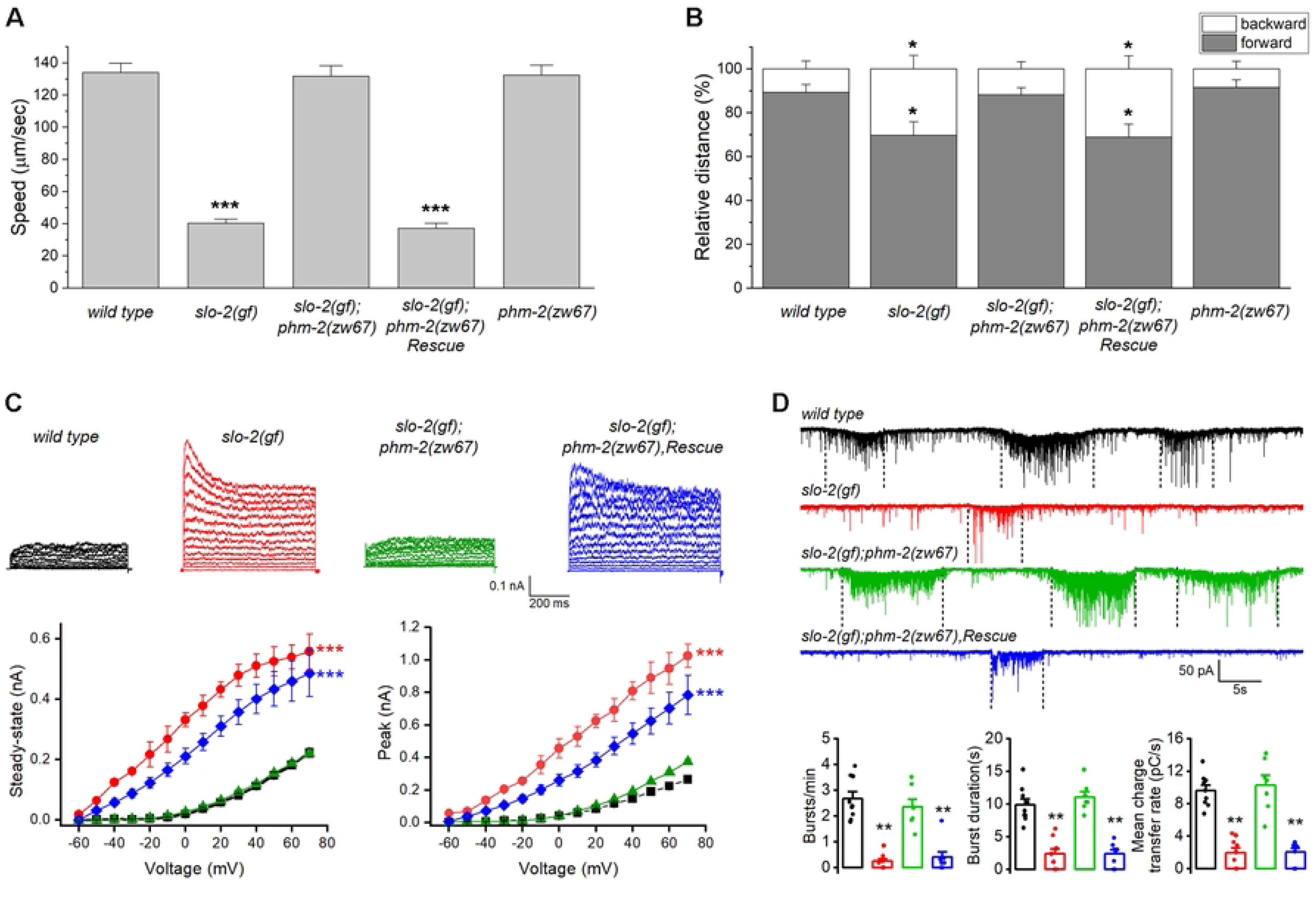
*phm-2* mutants suppress the lethargy of *slo-2(gf)*. (**A**) Comparison of average locomotion speed among various worm strains. (**B**) Comparison of the percentage of forward and backward locomotion among various worm strains. For both **A** and **B**, the sample sizes were 20 *wild type*, 18 *slo-2(gf)*, 17 *phm-2(zw67);slo-2(gf),* 15 rescue, and 19 *phm-2(zw67)*. The asterisk (*) indicates a statistically significant difference compared with wild type (* *p* < 0.05, *** *p* < 0.001, one-way *ANOVA* with Tukey’s post hoc tests). (**C**) *phm-2(zw67)* reverses the effects of *slo-2(gf)* on whole-cell currents in VA5 motor neuron, including larger amplitudes of peak and steady-state currents, and an apparent voltage-dependent inactivation, and these effects are rescued by expression of the wild-type *phm-2* in neurons. The sample sizes were 11 *wild type* (*wt*), 9 *slo-2(gf)*, 9 *phm-2(zw67);slo-2(gf),* and 9 rescue. The asterisk (*) indicates a significant difference compared with *wild type* (*** *p* < 0.001, two-way *ANOVA* with Tukey’s post hoc tests). (**D**) *phm-2(zw67)* suppresses inhibitory effects of *slo-2(gf)* on the frequency, duration, and mean charge transfer of postsynaptic current (PSC) bursts, and these effects are rescued by expression of the wild-type *phm-2* in neurons. Shown are representative traces of spontaneous PSCs and the comparison of PSC burst properties. The vertically dotted lines mark PSC bursts, which are defined as an apparent increase in PSC frequency accompanied by a baseline shift lasting > 3 sec. The sample sizes were 9 *wild type*, 9 *slo-2(gf)*, 7 *phm-2(zw65);slo-2(gf)*, and 8 rescue. The asterisk (*) indicates a statistically significant difference compared with *wild type* (** *p* < 0.01, one-way *ANOVA* with Tukey’s post hoc tests).

Expressing wild type *phm-2* in *slo-2(gf);phm-2(zw67)* double mutant fully reinstated the sluggish locomotion phenotype (**Figure 1A, B**), which confirms that the suppression of *slo-2(gf)* phenotype was indeed attributed to the mutation of *phm-*2. PHM-2 is an ortholog of human scaffold attachment factor B (SAFB) proteins (45). Like other SAFB proteins, PHM-2 has putative DNA-binding and RNA-binding domains, and arginine/glycine motifs. The mutation in *zw67* results in a premature stop at arginine 240 of PHM-2. We also analyzed the locomotion behaviors of the *phm-2(zw67)* mutant and found that *phm-2(zw67)* mutant has similar speed and forward/backward ratio as those of wild type worms (**Figure 1A, B**), suggesting that the suppression of *slo-2(gf)* phenotype by *phm-2(zw67)* mutant did not result from an additive effect. Thus, *phm-2* likely acts in the same genetic pathway to counteract *slo-2(gf)*.

To determine how *phm-2* mutant might alleviate the locomotion defect of *slo-2(gf)* worms, we recorded voltage-activated whole-cell currents from a representative ventral cord motor neuron (VA5) of different strains including wild type, *slo-2(gf)*, *slo-2(gf);phm-2(zw67)*, and *slo-2(gf);phm-2(zw67)* with *phm-2* rescued in neurons. Similar to what we reported previously (43, 46), VA5 whole-cell currents of *slo-2(gf)* worms were much bigger than those of wild type worms, with increases in both peak and sustained currents, and an apparent voltage-dependent inactivation that was not observed in wild type (**Figure 1C**). These characteristics of VA5 whole-cell currents of *slo-2(gf)* worms were completed abolished in *slo-2(gf);phm-2(zw67)* worms but could be restored in the double mutant by expressing wild-type *phm-2* in neurons (**Figure 1C**), suggesting that *phm-2* mutant has a direct effect to mitigate SLO-2(*gf*) function.

At *C. elegans* neuromuscular junctions, ventral cord motor neurons control body-wall muscle cells by producing postsynaptic current (PSC) bursts (47), and SLO-2 plays an important role in this function (31). To obtain further evidence that the suppression of *slo-2(gf)* phenotype by *phm-2* mutant was due to a dysfunction of SLO-2(*gf*) channel, we compared PSC bursts recorded from body-wall muscle cells of different strains. We found that PSC bursts in *slo-2(gf)* strain showed significant decreases in the frequency and strength compared with those in wild type worms (**Figure 1D**). These synaptic phenotypes of *slo-2(gf)* were not observed in the *slo-2(gf);phm-2(zw67)* strain but restored when wild-type *phm-2* was expressed in neurons of the double mutant (**Figure 1D**), suggesting that the activity of SLO-2*(gf)* channel was inhibited by the *phm-2* mutation. Together, our analyses of locomotion behaviors, neuronal whole-cell currents, and postsynaptic currents indicate that the action of SLO-2(*gf*) channel depends on PHM-2.

### PHM-2 is highly expressed in neurons and localized in the nucleus

Next, we examined the expression pattern of *phm-2* by expressing a promoter::GFP transcriptional reporter though an *in vivo* homologous recombination approach. Because reporters generated by this strategy incorporate larger genomic regions that are likely to contain all relevant cis-regulatory elements, they often better recapitulate endogenous gene expression (48). We cloned a 1.1-kb genomic fragment that includes part of the first common exon of all *phm-2* splice variants along with the upstream intron, and fused it to GFP. The resultant plasmid was then linearized and co-injected with a fosmid clone that contains part of *phm-2* coding region and 30-kb upstream sequence. *In vivo* homologous recombination between the plasmid and the fosmid is expected to result in a promoter::GFP transcriptional reporter that includes the genomic sequence upstream of *phm-2* in the fosmid. In transgenic worms, high GFP expression was observed in many neurons in the head and ventral nerve cord (**Figure 2A**). This expression pattern appears to be more restricted than that reported in a previous study, which observed ubiquitous *phm-2* expression using a 475-bp promoter fragment (45). The differences in cell-type-specific expression pattern observed with our *in vivo* recombination approach might be due to the existence of enhancer/repressor elements located in the distal upstream region of the *phm-2* locus.

**Figure 2.**
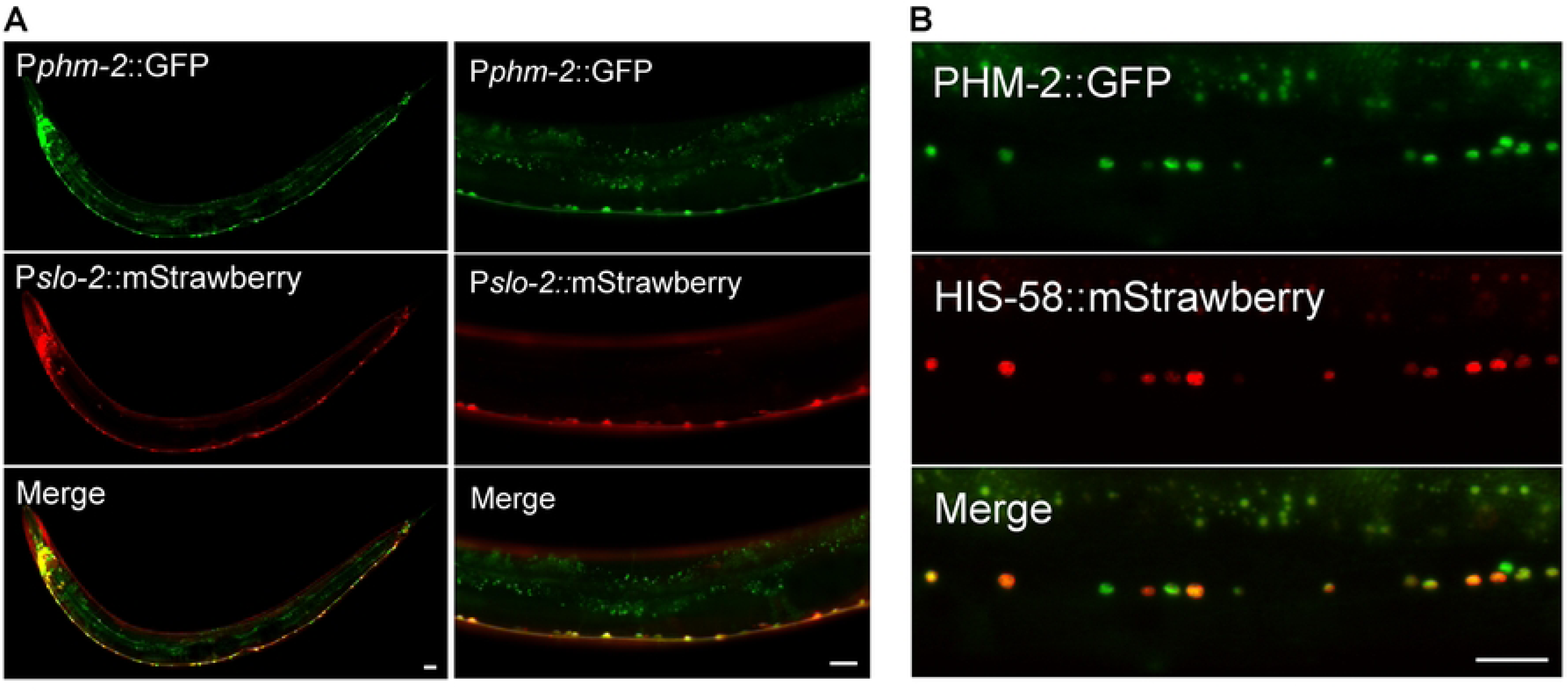
Expression and subcellular localization patterns of PHM-2. (**A**) *phm-2* is co-expressed with *slo-2* in many neurons in the nerve ring and along the ventral nerve cord. GFP and mStrawberry are expressed under the control of *phm-2* promoter (P*phm-2*) and *slo-2* promoter (P*slo-2*), respectively. (**B**) PHM-2 is localized in the nucleus of neurons. Shown are fluorescence images of a segment of the ventral nerve cord of a transgenic worm expressing a GFP-tagged PHM-2 and an mStrawberry-tagged HIS-58 (nucleus marker). Both fusion proteins are expressed under the control of the pan-neuronal *rab-3* promoter. Scale bar = 20 µm in all panels.

To determine whether the expression pattern of *phm-2* correlates with that of *slo-2*, we crossed the P*phm-2::gfp* reporter into a strain carrying P*slo-2::mStrawberry*, and examined the resulting GFP and mStrawberry signals. Co-expression of the two fluorescent markers was observed in many neurons in the head as well as in ventral nerve cord motor neurons (**Figure 2A**). Although some differences were noted between the GFP and mStrawberry patterns, both fluorescent proteins were strongly expressed in the same neuronal populations.

To determine the subcellular localization pattern of PHM-2, we cloned the longest *phm-2* splice variant and fused GFP at its C terminus. The PHM-2::GFP fusion protein was co-expressed with a nuclear marker HIS-58::mStrawberry in neurons. We found that the GFP signal overlapped with the mStrawberry signal (**Figure 2B**), consistent with the nuclear localization of SAFB proteins.

### PHM-2 regulates synaptic transmission through SLO-2

In wild-type worms, SLO-2 channels serve as the major carrier of the delayed outward currents in ventral cord motor neurons (31). To determine whether PHM-2 is required for native SLO-2 function in neurons, we recorded and compared VA5 whole-cell currents from wild type, *slo-2(lf)*, and *phm-2* mutant worms. Compared with wild type, VA5 whole-cell currents were greatly reduced in both *slo-2(lf)* and *phm-2(zw67)* mutants (**Figure 3A, B**). This mutant phenotype was confirmed in a new *phm-2* knockout allele, *zw93*, in which a stop codon was introduced after the amino acid residue alanine 495 of PHM-2 using the CRISPR/Cas9 approach. In *slo-2(lf);phm-2(zw93)* double mutant, VA5 whole-cell currents were not further reduced compared with those recorded from either *slo-2(lf)* or *phm-2* single mutants (**Figure 3A, B**), suggesting that PHM-2 contributes to motor neuron whole-cell currents through SLO-2. We also compared the resting membrane potential of VA5 between different strains, and found that it was less hyperpolarized in both *slo-2(lf)* and *phm-2* mutants than in wild type, and similar between *slo-2(lf);phm-2(zw93)* double mutant and the single mutants (**Figure 3C**), suggesting that PHM-2 plays a role in setting the resting membrane potential through SLO-2.

**Figure 3.**
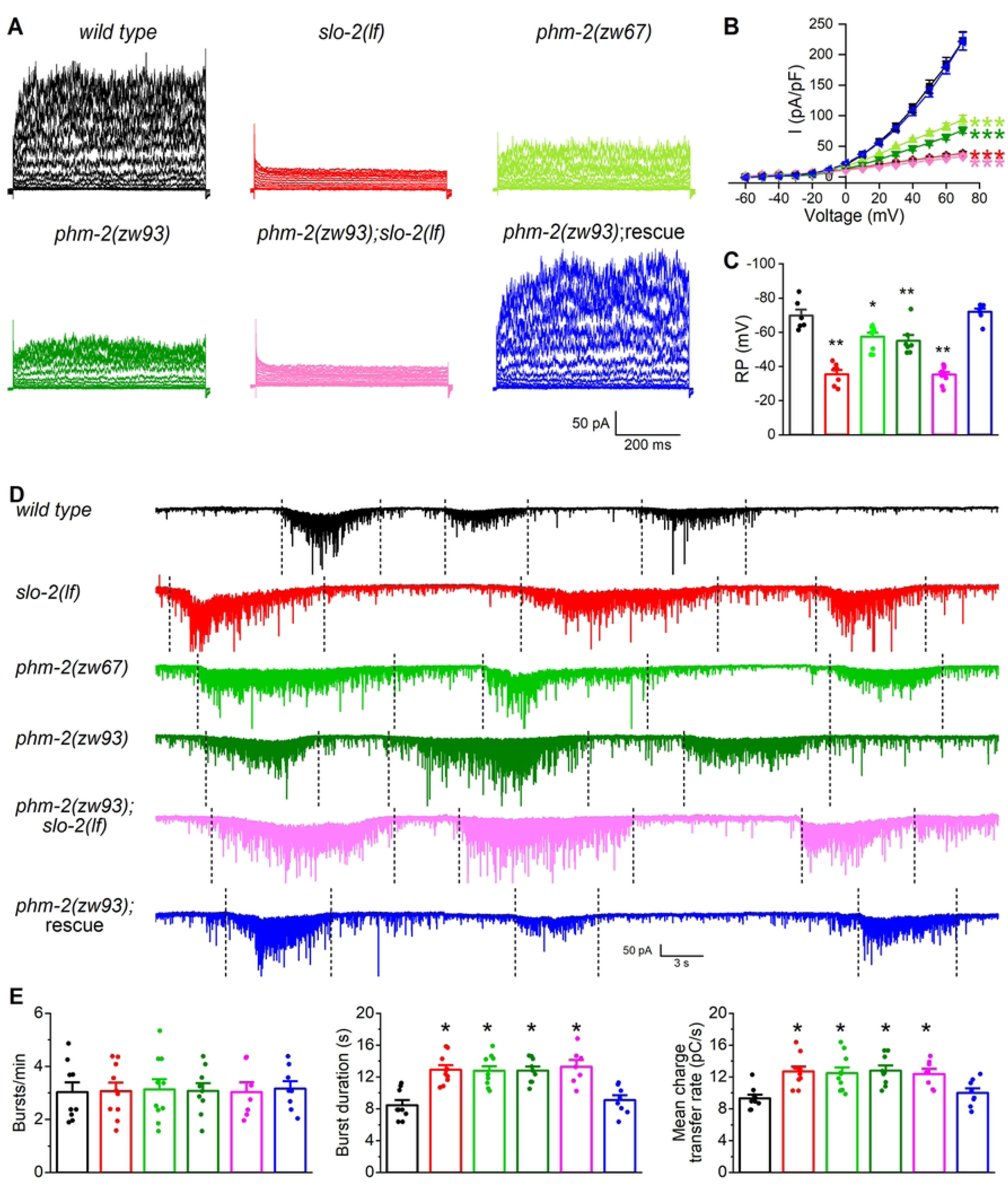
PHM-2 contributes to motor neuron whole-cell currents and regulates postsynaptic current (PSC) bursts through SLO-2. (**A**) Representative VA5 whole-cell current traces. (**B**) Current (*I*) - voltage relationships of the whole-cell currents. Sample sizes were 11 *wild type*, 9 *slo-2(lf)*, 9 *phm-2(zw67)*, 12 *phm-2(zw93)*, 12 *phm-2(zw93);slo-2(lf),* and 12 *phm-2(zw93)* rescue. (**C**) Resting membrane potentials of VA5. Sample sizes were 6 *wild type*, 7 *slo-2(lf)*, 9 *phm-2(zw67)*, 7 *phm-2(zw93)*, 10 *phm-2(zw93);slo-2(lf),* and 7 *phm-2(zw93)* rescue. (**D**) Representative traces of spontaneous PSCs with PSC bursts marked by vertically dotted lines. (**E**) Comparisons of PSC burst properties. Sample sizes were 9 *wild type*, 10 *slo-2(lf)*, 10 *phm-2(zw67)*, 9 *phm-2(zw93)*, 7 *phm-2(zw93);slo-2(lf),* and 8 *phm-2(zw93)* rescue. All values are shown as mean ± SE. The asterisks indicate statistically significant differences (\**p* < 0.05, \*\**p* < 0.01, \*\*\**p* < 0.001) compared with *wild type* based on either two-way (**B**) or one-way (**C** and **E**) ANOVA with Tukey’s post hoc tests.

SLO-2 serves as a negative regulator of neurotransmitter release at the neuromuscular junctions (31), and *slo-2(lf)* mutants exhibit significantly increased duration and charge transfer rate of PSC bursts. To determine whether PHM-2 is also required for this function of SLO-2, we recorded PSC bursts from body-wall muscle cells of wild type and various mutant strains. Similar to *slo-2(lf)* mutant, the *phm-2* mutants showed increased duration and charge transfer rate of PSC bursts without a change in burst frequency. These synaptic phenotypes of *phm-2* mutants were not additive with those of *slo-2(lf)* and were rescued by expressing wild-type *phm-2* specifically in neurons (**Figure 3D, E**), indicating that PHM-2 regulates neurotransmitter release through presynaptic SLO-2.

### PHM-2 is required for SLO-2 expression

As an RNA/DNA binding protein, PHM-2 might regulate neuronal function through an effect on SLO-2 expression. To test this possibility, an ideal approach would be to compare native SLO-2 protein levels between wild type and *phm-2* mutant worms using an antibody against SLO-2. However, no commercial SLO-2 antibodies were available, and our effort to generate a custom SLO-2 antibody was unsuccessful. Therefore, we examined SLO-2 expression using a SLO-2::GFP fusion protein. To maintain identical transgene dosage, we crossed a highly stable transgene (> 99% penetrance) expressing SLO-2::GFP in neurons from an existing wild-type strain into two *phm-2* mutants, and quantified the GFP epifluorescence in a segment of the ventral nerve cord. We found that GFP epifluorescence in neurons was greatly reduced in both *phm-2* mutants compared with wild-type worms (**Figure 4A, B**), suggesting that PHM-2 is required for proper SLO-2 expression. To determine whether the decrease in SLO-2::GFP in *phm-2* mutants reflects a global reduction of protein expression, we compared SLO-1::GFP expression between wild type and *phm-2(zw93)* worms. SLO-1 is a voltage- and calcium-activated potassium channel and a paralog of SLO-2 in worms. We found that SLO-1::GFP expression in *phm-2(zw93)* mutants remained unchanged relative to wild type (**Figure 4C**). Thus, knockout of *phm-2* does not cause a general reduction in protein expression.

**Figure 4.**
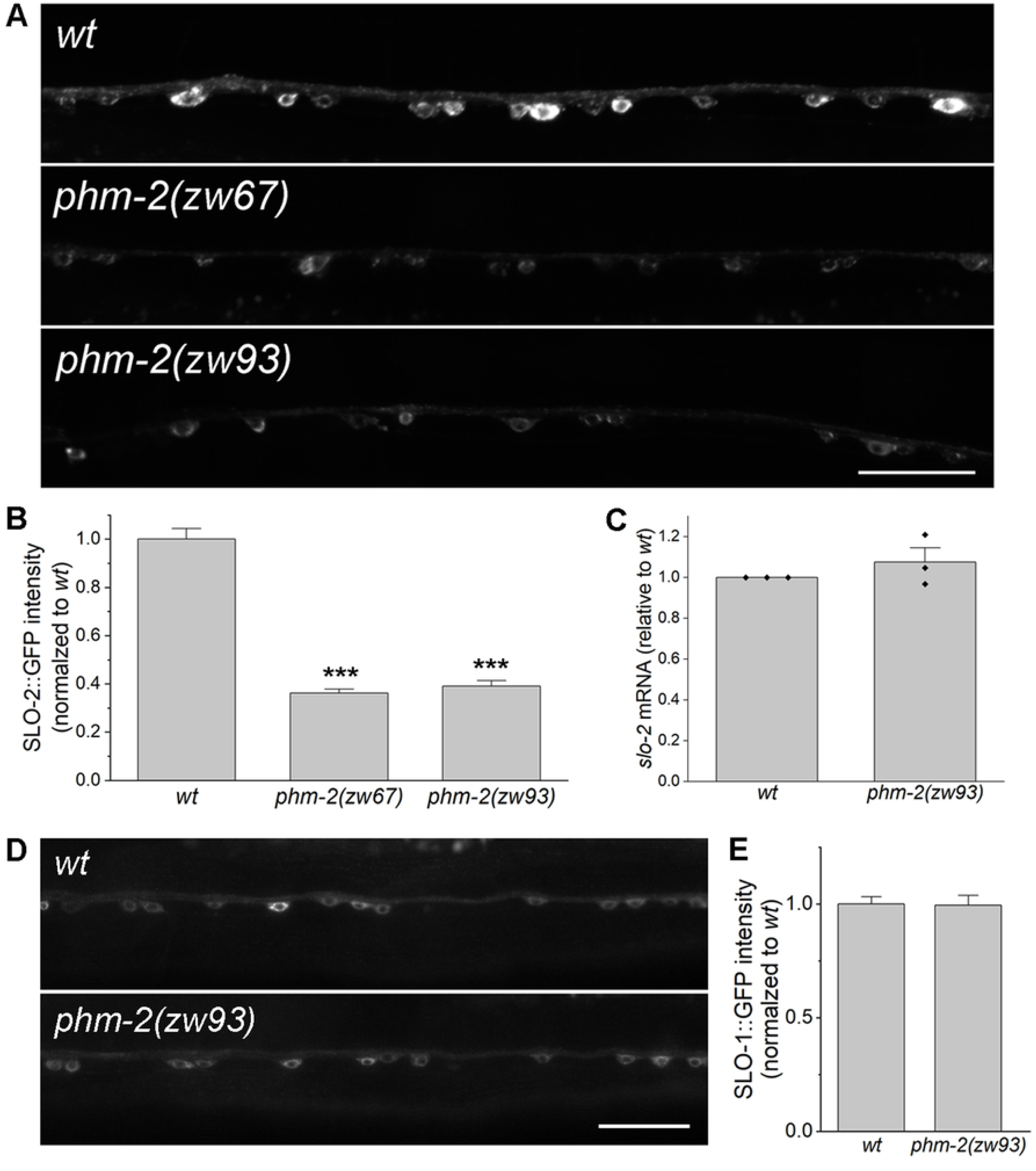
PHM-2 regulates SLO-2 expression post-transcriptionally. (**A**) Representative images of SLO-2::GFP expression in the ventral nerve cord from wild type (*wt*) and *phm-2* mutants. Scale bar = 20 µm. (**B**) Comparison of SLO-2::GFP signal between *wt* and *phm-2* mutants. The sample sizes were 28 *wt*, 22 *phm-2(zw67)*, and 25 *phm-2(zw93)*. The asterisk (*) indicates a significant difference compared with *wt* (*** *p* < 0.001, one-way *ANOVA* with Tukey’s post hoc tests). (**C**) Comparison of *slo-2* mRNA level between *wt* and *phm-2(zw93)* mutants. *slo-2* mRNA level in the mutant was normalized to *wt* = 1. A one-sample t-test indicated no significant difference from *wt* (*p* = 0.4045). (**D**) Representative images of SLO-1::GFP expression in ventral nerve cord from *wt* and *phm-2*(*zw93*) mutants. Scale bar = 20 µm. ***E***, Comparison of SLO-1::GFP signal between *wt* and *phm-2(zw93)* mutants. SLO-1::GFP signal was normalized to *wt*. A two-sample (unpaired) Student’s t-test indicated no significant difference between the two groups (*p* = 0.9387). The sample sizes were 16 *wt* and 19 *phm-2(zw93)*.

PHM-2 might regulate SLO-2 expression through transcriptional or post-transcriptional mechanisms. To distinguish these possibilities, we quantified *slo-2* mRNA levels in wild type and the *phm-2(zw93)* mutant using our RNA-Seq data. We found that *slo-2* mRNA levels were the same between wild type and *phm-2(zw93)* mutant (**Figure 4D**), suggesting that PHM-2 regulates SLO-2 expression post-transcriptionally.

### Decreased SLO-2 expression is not caused by an innate immune response in phm-2(lf) mutant

A previous study showed that *phm-2* mutants have an abnormal pharyngeal grinder that allows live bacteria to accumulate in the intestine, which triggers innate immune responses that result in a scrawny body morphology and delayed reproductive aging. These mutant phenotypes can be abrogated by feeding the mutants UV-killed *E. coli* (45). To examine whether the effect of *phm-2* mutants on SLO-2 expression might involve an immune response, we analyzed SLO-2::GFP expression in wild-type and *phm-2(zw93)* worms fed either live or UV-killed *E. coli*. We found that SLO-2::GFP expression was unchanged between the two diets in both wild type and *phm-2(zw93)* mutants (**Figure 5A**). We also recorded VA5 whole-cell currents from these worms and found no significant diet-dependent differences within each genotype (**Figure 5B**). These observations suggest that the effect of *phm-2* mutation on SLO-2 expression was not due to an immune response.

**Figure 5.**
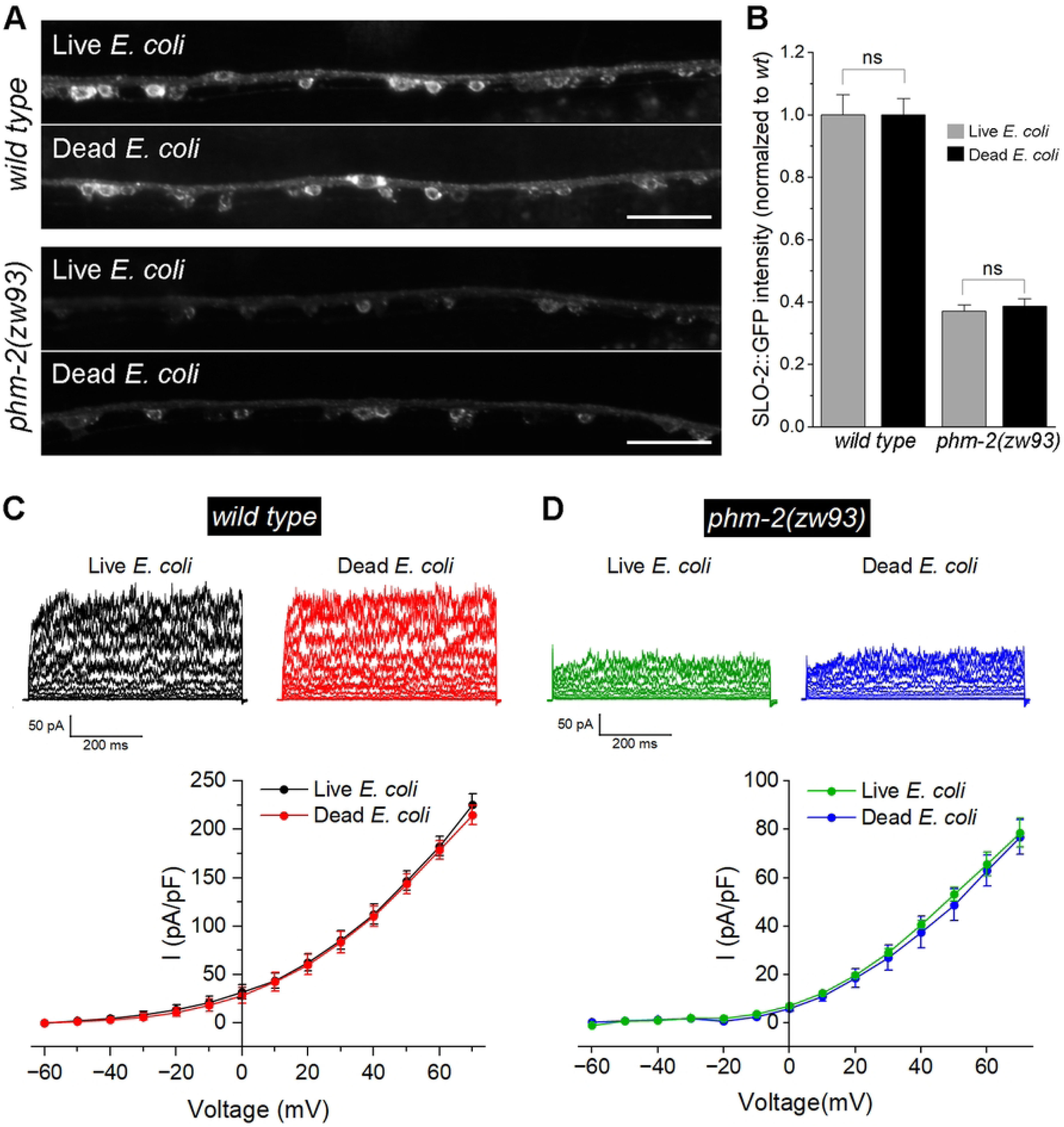
The innate immune response in *phm-2* mutants does not cause a reduction of SLO-2 expression in neurons. (**A**) Representative images of SLO-2::GFP expression in the ventral nerve cord of wild type and *phm-2(zw93)* mutants grown on either live or dead *E. coli.* Scale bar = 20 µm. (**B**) Comparison of SLO-2::GFP signals in worms grown on live versus dead *E. coli*. The sample sizes were 19 *wt* (live *E. coli*), 21 *wt* (dead *E. coli*), 19 *phm-2(zw93)* (live *E. coli*), and 18 *phm-2(zw93)* (dead *E. coli*). Statistical significance between live and dead *E. coli* groups within each genotype was evaluated using a two-sample (unpaired) Student’s t-test. Bonferroni correction was applied for two comparisons (adjusted α = 0.025). No statistically significant differences were detected (*p* > 0.05). (**C**) Comparison of VA5 whole-cell currents in wild-type worms grown on live versus dead *E. coli*. The sample sizes were 10 grown on live *E. coli*, 10 grown on dead *E. coli*. (**D**) Comparison of VA5 whole-cell currents in *phm-2(zw93)* mutants grown on live versus dead *E. coli*. The sample sizes were 7 grown on live *E. coli*, 10 grown on dead *E. coli*. In both **C** and **D**, no statistically significant differences were detected between the groups (*p* > 0.05, two-way *ANOVA* with Tukey’s post hoc tests).

### PHM-2 acts with HRPU-2 to control SLO-2 expression

Our previous study showed that SLO-2 expression is regulated by HRPU-2, a homologue of mammalian heterogeneous nuclear ribonucleoprotein U (hnRNP U) (43). Because HRPU-2 regulates SLO-2 expression through a post-transcriptional mechanism, we asked whether PHM-2 and HRPU-2 might function together to regulate SLO-2. To test this, we generated a *phm-2(zw93);hrpu-2(zw97)* double mutant and compared SLO-2::GFP expression between the double mutant and each single mutant. The *hrpu-2(zw97)* mutant allele was created by introducing a stop codon after the amino acid residue glutamate 194 of HRPU-2 using the CRISPR/Cas9 approach. We found that the SLO-2::GFP intensity in neurons was similar among the double and the single mutants (**Figure 6A, B**). These results suggest that PHM-2 and HRPU-2 likely act in the same pathway to control SLO-2 expression.

**Figure 6.**
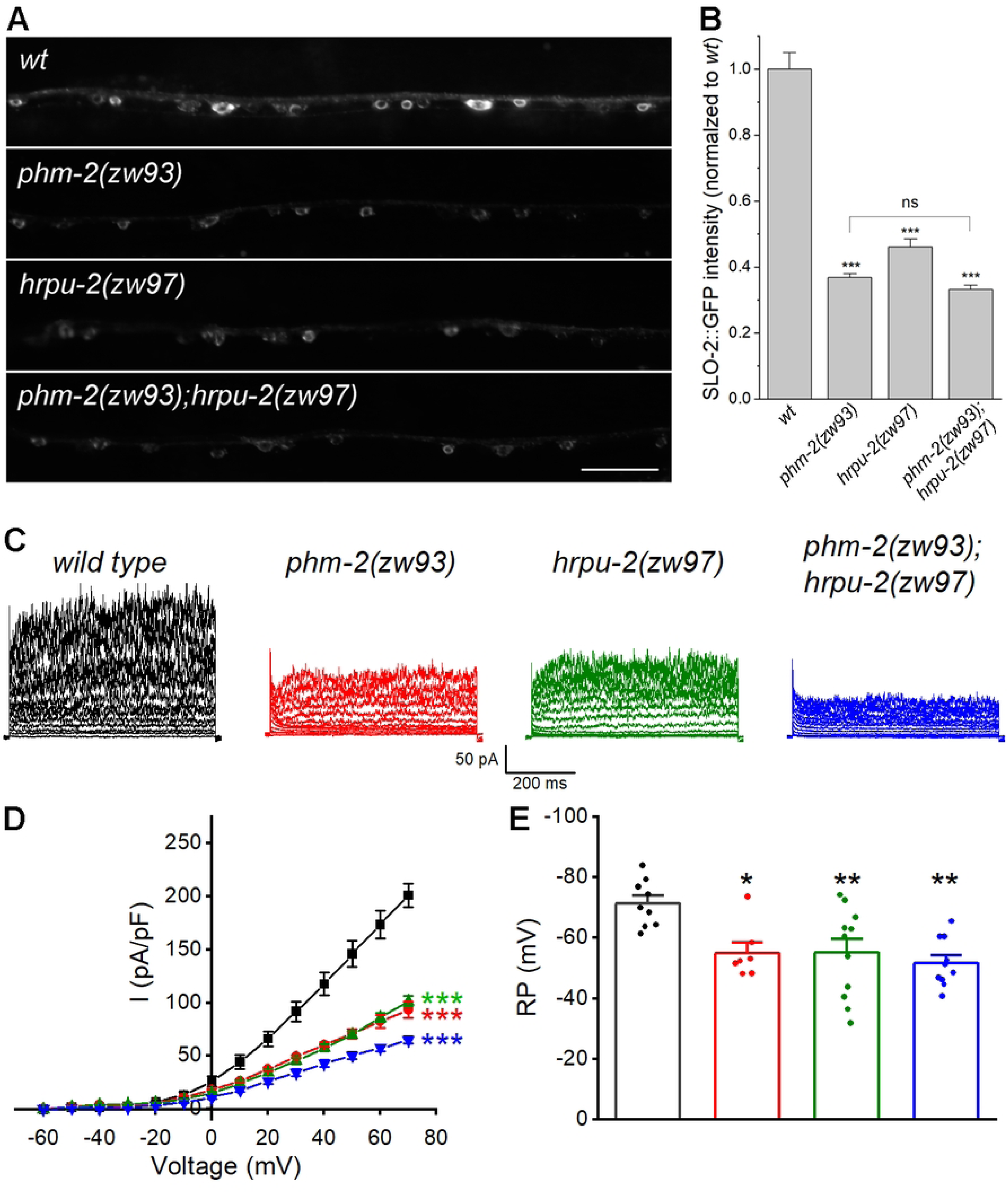
PHM-2 acts with HRPU-2 to control SLO-2 expression. (**A**) Representative images of SLO-2::GFP expression in the ventral nerve cord from *wild type* (*wt*) and the single and double mutants. Scale bar = 20 µm. (**B**) Comparison of SLO-2::GFP signal between *wt* and the single and double mutants. The sample sizes were 24 *wt*, 25 *phm-2(zw93)*, 22 *hrpu-2(zw97)*, and 25 *phm-2(zw93)*;*hrpu-2(zw97)*. The asterisk (*) indicates a significant difference compared with *wt* (*** *p* < 0.001, one-way *ANOVA* with Tukey’s post hoc tests). (**C**) Representative VA5 whole-cell current traces. (**D**) Current (*I*) - voltage relationships of the whole-cell currents. Values are shown as mean ± SE. The asterisks indicate statistically significant differences compared with *wt* (\*\*\**p* < 0.001, two-way ANOVA with Tukey’s post hoc tests). Sample sizes were 9 *wt*, 7 *phm-2(zw93)*, 11 *hrpu-2(zw97)*, and 10 *phm-2(zw93)*;*hrpu-2(zw97)*. (**E**) Resting membrane potentials of VA5. Data are shown as mean ± SE. The asterisks indicate statistically significant differences compared with *wt* (\**p* < 0.05, \*\**p* < 0.01, one-way ANOVA with Tukey’s post hoc tests). Sample sizes were 9 *wt*, 7 *phm-2(zw93)*, 11 *hrpu-2(zw97)*, and 10 *phm-2(zw93)*;*hrpu-2(zw97)*.

To further examine the relationship between PHM-2 and HRPU-2 with respect to SLO-2 function, we compared VA5 whole-cell currents in the *phm-2(zw93);hrpu-2(zw97)* double mutant and those in the single mutants. If PHM-2 and HRPU-2 regulate SLO-2 independently, we would expect to see very little currents in the double mutant. Instead, VA5 whole-cell currents in the double mutant were only slightly smaller than those in either single mutant (**Figure 6C, D**), indicating that the current reductions in the single mutants largely overlap. In addition, VA5 resting membrane potentials were comparable between the single and double mutants (**Figure 6E**). Taken together, these results suggest that PHM-2 and HRPU-2 contribute to neuronal function by acting together on SLO-2 expression.

## Discussion

In this study, we demonstrate that PHM-2, the sole SAFB protein in *C. elegans*, plays an important role in synaptic transmission through the SLO-2 potassium channel. Our conclusion is supported by multiple lines of evidence at the behavioral, cellular, and molecular levels: first, *phm-2* mutants suppress the sluggish phenotype of *slo-2(gf)* worms; second, PHM-2 contributes to neuronal whole-cell currents and post-synaptic currents through SLO-2; third, PHM-2 controls SLO-2 expression through a post-transcriptional mechanism. In addition, we found that PHM-2 functions with another RBP, HRPU-2/hnRNP U, to regulate SLO-2. These findings define a novel neuronal function of a SAFB protein, and may help our understanding of SAFB proteins and their targets in physiology and disease.

We previously showed that HRPU-2/hnRNP U regulates SLO-2 expression in *C. elegans* neurons (43). Our new results suggest that PHM-2 has a similar role and may act together with HRPU-2 to regulate SLO-2. In mammals, SAFB1 and hnRNP U were initially characterized based on their capability of binding to matrix/scaffold attachment region (S/MAR) of DNA elements (49, 50). Subsequent studies suggest that these proteins may interact to regulate specific cellular processes. For example, in a comprehensive analysis of the long non-coding RNA (lncRNA)-bound proteome using ChIRP-MS, both SAFB1 and hnRNP U were identified among proteins that bind Xist (51), an lncRNA required for X-chromosome inactivation in female cells to achieve dosage compensation males (52). In addition, hnRNP

U was efficiently pulled down along with several other hnRNPs as SAFB1-interacting proteins from Hela cell nuclear extracts (12). In humans, mutations of the Slo2.2/Slack channel are strongly linked with epilepsies and intellectual disability (53-58), and deficiencies of hnRNP U are similarly associated with epileptic encephalopathies and intellectual disability (59-64). Notably, a clinical study reported that SAFB1 and SAFB2 might be responsible for epilepsy and mental retardation in a patient (65). The common clinical presentations observed in patients with mutations in these genes, together with our findings in worms, suggest that these molecules may interact to perform critical neuronal functions.

A previous study showed that *phm-2* mutants have a pharynx grinder defect, which leads to accumulation of live bacteria in the intestine and activation of innate immune responses, including a bacterial avoidance behavior. As a result, *phm-2* mutants exhibit a scrawny body morphology and delayed somatic and reproductive aging (45). These phenotypes of *phm-2* mutants do not appear to be related to either SLO-2 or HRPU-2 because both *slo-2(lf)* and *hrpu-2* mutants are morphologically similar to wild-type worms (43). In addition, while the morphological phenotypes of *phm-2* mutants are abrogated when they are raised on a UV-killed *E. coli* diet (45), the decreased SLO-2 expression and whole-cell currents in neurons of *phm-2* mutants remain unchanged compared with those fed a normal *E. coli* diet. These observations suggest that PHM-2 has multiple physiological functions, which is consistent with its broad expression pattern. In mammals, SAFB proteins are widely expressed, with high expression observed in the developing and mature brain (25). Immunoblot data also show high expression of SAFB1 and SAFB2 in the immune system and in hormonally regulated organs such as uterus and ovary (66). It would be interesting to know whether SAFB proteins may have conserved roles in the immune and reproductive systems.

Slo2 channels are a family of evolutionarily conserved potassium channels. They are widely expressed in the nervous system and have important roles in shaping neuronal activity. Like many other ion channels, the activity of Slo2 channels is regulated by a variety of signaling molecules, such as G-protein-coupled receptors, protein kinases, cyclic AMP, phosphatidylinositol 4,5-biphosphate (PIP2), nicotinamide adenine dinucleotide NAD+, and others (67). Interestingly, the physiological functions of Slo2 channels may also be regulated by specific RBPs, either directly or indirectly. In mice, the Fragile X Mental Retardation Protein (FMRP) interacts directly with the cytoplasmic C-terminus of the Slo2.2/Slack subunit and potently activates the channel (68). In *C. elegans* neurons, HRPU-2 binds to *slo-2* mRNA and regulates its expression post-transcriptionally (43), and PHM-2 plays a similar role to HRPU-2 and may interact with it to regulate SLO-2 expression. In addition, SLO-2 function is indirectly modulated by ADR-1 (46), an RBP that regulates adenosine-to-inosine RNA editing (69, 70). ADR-1 facilitates expression of SCYL-1, an evolutionarily conserved regulator of Slo2 channels, by promoting editing of a single adenosine at the 3’-UTR of *scyl-1* transcripts (46). These findings suggest that neuronal Slo2 channels might be under the control of a complex network of RBPs; however, the physiological significance of such regulation remains to be determined.

Our results show that, similar to HRPU-2, PHM-2 regulates *slo-2* expression post-transcriptionally. Several mechanisms could be responsible for this regulation, including alternative splicing, nuclear editing, mRNA trafficking, mRNA stability, and translation. In our experiments comparing SLO-2 expression between *phm-2* mutants and wild-type worms, a *slo-2* cDNA was used to express the SLO-2::GFP fusion in neurons, and the SLO-2(*gf*) channel that causes the sluggish phenotype suppressed by *phm-2* mutants is also encoded by a *slo-2* cDNA with a few modifications. Thus, alternative splicing does not appear to be involved in the regulation of SLO-2 by PHM-2. In addition, no RNA editing events are detected in *slo-2* mRNA (46), and *slo-2* mRNA levels are similar between wild type and *phm-2* mutants. Therefore, it appears that, like HRPU-2 (43), PHM-2 may regulate SLO-2 expression through either *slo-2* mRNA transport or translation. Nevertheless, apart from their common role in regulating SLO-2, HRPU-2 and PHM-2 likely have distinct functions since the phenotypes of *phm-2* and *hrpu-2* mutants are very different (43, 45).

In the brain, RBPs are essential for maintaining and controlling neuronal excitability (71). The interaction between RBPs and their targets may link neuronal activity to gene expression that is critical for synaptic plasticity. Indeed, many studies have implicated RBPs in neuronal activity-dependent regulation of gene expression through various post-transcriptional mechanisms, including alternative splicing (72-74), RNA trafficking (75-77), and translation (78-80). Given that Slo2 channels play major roles in shaping neuronal firing patterns, their regulation by RBPs may be an important mechanism by which the nervous system maintains its homeostasis in response to changes in neuronal activity.

## Materials and Methods

### C. elegans culture and strains

*C. elegans* were cultured on nematode growth medium (NGM) plates seeded with a layer of *Escherichia coli* OP50 at 22°C inside an environmental incubator. The following strains were used in this study: wild type (Bristol N2). LY101: *slo-2(nf101)*. ZW083: *zwIs101[Pslo-1::slo-1::GFP]*. ZW860: *zwIs[Pslo-1::slo-2(gf), Pmyo-2::yfp]*. ZW863: *zwIs[Pslo-1::slo-2(gf), Pmyo-2::yfp]; phm-2(zw67)*. ZW975: *zwEx221[Prab-3::slo-2::GFP]*; *phm-2(zw67).* ZW996: *phm-2(zw67)*. ZW1051: *zwEx221[Prab-3::slo-2::GFP]; hrpu-2(zw97)*. ZW1049: *zwEx221[Prab-3::slo-2::GFP]*. ZW1070: *phm-2(zw93)*. ZW1357: *phm-2(zw93)*;*slo-2(nf101).* ZW1365: *zwEx401[Prab-3::phm-2, Pmyo-2:: mStrawberry]; phm-2(zw93)*. ZW1387: *zwEx247[Pslo-2::mStrawberry, lin-15(+)]; zwEx403[Pphm-2::GFP, fosmid WRM064cE02, lin-15(+)]; lin-15(n765).* ZW1395: *hrpu-2(zw97).* ZW1404: *zwEx221[Prab-3::slo-2::GFP]*; *phm-2(zw93).* ZW1609: *zwEx402[Prab-3::phm-2::GFP, Prab-3::His-58::mStrawberry, lin-15(+)]; lin-15(n765)*. BJC201: *zwIs101[Pslo-1::slo-1::GFP]*; *phm-2(zw93).* BJC272: *phm-2(zw93); hrpu-2(zw97).* BJC273: *zwEx221[Prab-3::slo-2::GFP]; phm-2(zw93);hrpu-2(zw97)*.

### Mutant screening and mapping

Mutant screen was performed with a *C. elegans* strain with an integrated transgenic array expressing P*slo-1::slo-2(gf)* and P*myo-2::yfp* in the N2 Bristol background. L4-stage hermaphrodite worms were immersed in M9 buffer containing 50 mM ethyl methanesulfonate (EMS) for 4 hours at room temperature, and transferred to NGM plates after washing with M9 buffer 5 time. F1 progeny of EMS-treated worms were placed on NGM plates (∼10 worms/plate), and the F2 worms were screened for mutants that moved better than the original *slo-2(gf)* worms. The isolated mutants were subjected to whole-genome sequencing, and SNP-based genetic mapping was performed to determine the rough chromosome location of the mutations. One of the mutants, *zw67*, was mapped to *F32B4.4 (phm-2)* and confirmed by subsequent rescue experiments.

### Analysis of locomotion behavior

Locomotion behavior was analyzed using an automated worm tracking and analysis system (81). Briefly, a single young adult hermaphrodite was placed on an NGM plate without food. After ∼30 sec recovery time from the transfer, snapshots of the worm were taken at 15 frames per sec for 30 s using an IMAGINGSOURCE camera (DMK37BUX273) mounted on a stereomicroscope (SMZ800, Nikon, Tokyo, Japan). The worm was constantly kept in the center of the view field with a motorized microscope stage (OptiScanTM ES111, Prior Scientific, Inc., Rockland, MA, USA). Both the camera and the motorized stage were controlled by a custom program running in MATLAB (The MathWorks, Inc., Natick, MA).

### Generation of phm-2 and hrpu-2 knockout strains

The CRISPR/Cas9 approach was used to create *phm-2* and *hrpu-2* knockout strain. To generate *phm-2* knockout, a guide RNA sequence 5’- GAGAAGCATGTTGCTGAGG was inserted into pDD162 (P*eft-3::Cas9* + Empty sgRNA; Addgene #47549). The resultant plasmid was injected into wild type worms along with a repair primer (5’-AAACTGGCTCGGGAGAAGCATGTTGCTTAATGAAGGCGGCGAGCACAATGAGCACTTCCC) and Pmyo-2::mStrawberry (*wp1613*) as the transgenic marker. The *phm-2* knockout worms were identified by PCR using primers 5’- GGGAGAAGCATGTTGCTTAATGA (forward) and 5’-GGAGATTGGAGGATTAGCGGA (reverse). The knockout worms were confirmed by Sanger sequencing. The *hrpu-2* knockout was generated using the same approach. The guide RNA sequence for hrpu-2 was 5’-TGGATCATGATGATGAAGG, and the repair primer sequence was: 5’-TGATGAATTAATGGATCATGATGATGAATAACTGGAGGGCATGATGAACATGAAGAAGAT Knockout worms were identified by PCR using primers 5’- TTAATGGATCATGATGATGAATAACT (forward) and 5’-TGTGACCGATGTCCAGTAATCC (reverse).

### Analysis of expression pattern and subcellular localization

The expression pattern of *phm-2* was assessed by an in vivo recombination approach. Specifically, a 1.1 kb fragment immediately upstream of phm-2 initiation site was cloned and fused to GFP using the primers 5’-TTTGGTACCAAGCCAGAAGAAATTCCACACAA (forward) and 5’- ATAACCGGTCCGTATTGGCTCGCGAGT (reverse). The resultant plasmid (P*phm-2::gfp, wp1888*) was linearized and co-injected with a linearized (fosmid WRM064cE02), which contains 30 kb of *phm-2* upstream sequence and part of its coding region, into the *lin-15(n765)* strain along with a *lin-15* rescue plasmid to serve as a transformation marker. Subcellular localization of PHM-2 was determined by fusing GFP to its carboxyl terminus and expressing the fusion protein under the control of P*rab-3* (P*rab-3::phm-2::gfp*, *wp1965*). Primers for cloning *phm-2* cDNA are 5’-TTTGGTACCATGCCGTTGGAAAGCGGAAAA (forward) and 5’- AATACCGGTCAATAATTTCCGCGATAATTTCCATA (reverse). A plasmid harboring P*rab-3::his-58::mStrawberry* (*wp1749*) was used to serve as a nucleus marker. The plasmids were injected into the *lin-15(n765)* strain along with a *lin-15* rescue plasmid to serve as a transformation marker. To determine whether *phm-2* is co-expressed with *slo-2*, the P*phm-2::gfp* transgene was crossed into an existing strain expressing P*slo-2::mStrawberry*. Images of transgenic worms were taken with a digital CMOS camera (Hamamatsu, C11440-22CU) mounted on a Nikon TE2000-U inverted microscope equipped with EGFP/FITC and mCherry/Texas Red filter sets (49002 and 49008, Chroma Technology Corporation, Rockingham, VT, USA).

### RNA-seq and data analysis

Total RNA was extracted from young adult-stage worms using TRIzol Reagent (Invitrogen) and treated with TURBO DNase (Ambion). RNA-seq was performed by Novogene Corp. Sacramento, CA. Raw reads ware filtered using Trim Galore software (http://www.bioinformatics.babraham.ac.uk/projects/trim_galore/) to remove reads containing adapters or reads of low quality. The filtered reads were mapped to *C. elegans* genome (ce11) using TopHat2 (Kim et al., 2013). The gene expression level is estimated by counting the reads that map to exons.

### Quantification of SLO-2::GFP and SLO-1::GFP fluorescence intensity

Young adult worms expressing P*rab-3::slo-2::gfp* or P*slo-1::slo-1::gfp* were immobilized in M9 solution containing 1mM azide. Images of the ventral nerve cords posterior to the vulva were obtained using the Hamamatsu digital CMOS camera with an identical exposure time for each group. The ImageJ software was used to extract straightened ventral cord images and to quantify fluorescence intensity. For each image, SGFP intensity was calculated by subtracting the minimum intensity (background fluorescence) from the average intensity.

### Electrophysiology

Adult hermaphrodites were used in all electrophysiological experiments. Worms were immobilized and dissected as described previously (47). Borosilicate glass pipettes were used as electrodes for recording whole-cell currents. Pipette tip resistance for recording muscle cell currents was 3-5 MΩ whereas that for recording motor neuron currents was ∼20 MΩ. The dissected worm preparation was treated briefly with collagenase and perfused with the extracellular solution for 5 to 10-fold of bath volume. Classical whole-cell configuration was obtained by applying a negative pressure to the recording pipette. Current- and voltage-clamp experiments were performed with a Multiclamp 700B amplifier (Molecular Devices, Sunnyvale, CA, USA) and the Clampex software (version 10, Molecular Devices). Data were sampled at a rate of 10 kHz after filtering at 2 kHz. Spontaneous membrane potential changes were recorded using the current-clamp technique without current injection. VA5 motor neuron whole-cell currents were recorded by applying a series of voltage steps (−60 to +70 mV at 10-mV intervals, 600 ms pulse duration) from a holding potential of −60 mV. The bath solution contained (in mM) 140 NaCl, 5 KCl, 5 CaCl_2_, 5 MgCl_2_, 11 dextrose and 5 HEPES (pH 7.2). The pipette solution contained (in mM) 120 KCl, 20 KOH, 5 Tris, 0.25 CaCl_2_, 4 MgCl_2_, 36 sucrose, 5 EGTA, and 4 Na_2_ATP (pH 7.2). Spontaneous PSCs were recorded from body-wall muscle cells at a holding potential of -60 mV. The recording solutions were the same as those for neuronal whole-cell currents recording, except that the 113.2 KCl in the pipette solution was substituted by K^+^ gluconate.

### Data Analyses for Electrophysiology

Amplitudes of whole-cell currents in response to voltage steps were determined from the mean current during the last 100 ms of the 600-ms voltage pulses using the Clampfit software. The duration and charge transfer of PSC bursts were quantified with Clampfit software (version 11, Molecular Devices) as previously described (47). The frequency of PSC bursts was manually counted. Statistical comparisons were performed with Origin Pro 2021 (OriginLab Corporation, Northampton, MA) using either *ANOVA* or unpaired *t*-test as specified in figure legends. *p* < 0.05 is considered to be statistically significant. The sample size (*n*) equals the number of cells or membrane patches analyzed. All values are shown as mean ± SE and data graphing was done with Origin Pro 2021.

## Acknowledgements

This work was supported by National Institute of Health (R35GM139620 to B.C). Some strains were provided by the CGC, which is funded by NIH Office of Research Infrastructure Programs (P40 OD010440).

